# Combining *δ*^13^C and *δ*^15^N from bone and dentine in marine mammal palaeoecological research: insights from toothed whales

**DOI:** 10.1101/2021.12.13.472362

**Authors:** Alba Rey-Iglesia, Tess Wilson, Jennifer Routledge, Mikkel Skovrind, Eva Garde, Mads Peter Heide-Jørgensen, Paul Szpak, Eline D. Lorenzen

## Abstract

Stable carbon (*δ*^13^C) and nitrogen (*δ*^15^N) isotope compositions of bone and dentine collagen extracted from museum specimens have been widely used to study the paleoecology of past populations. Due to possible systematic differences in stable isotope values between bone and dentine, dentine values need to be transformed into bone-collagen equivalent using a correction factor to allow comparisons between the two collagen sources. Here, we provide correction factors to transform dentine *δ^13^C* and *δ*^15^N values into bone-collagen equivalent for two toothed whales: narwhal and beluga.

We sampled bone and tooth dentine from the skulls of 11 narwhals and 26 belugas. In narwhals, dentine was sampled from tusk and embedded tooth; in beluga, dentine was sampled from tooth. *δ*^13^C and *δ*^15^N were measured and intraindividual bone and dentine isotopic compositions were used to calculate correction factors for each species.

We detected differences in *δ*^13^C and *δ*^15^N. In narwhals, we found (i) lower average *δ*^13^C and *δ*^15^N in bone compared with dentine; (ii) no difference in dentine *δ*^13^C between tusk and embedded tooth. For belugas, we also detected lower *δ*^13^C and *δ*^15^N in bone compared with tooth dentine. The correction factors provided by the study enable the combined analysis of stable isotope data from bone and dentine in these species.

## Introduction

Stable carbon (*δ*^13^C) and nitrogen (*δ*^15^N) isotope compositions of bone and dentine collagen extracted from subfossil specimens have been used to study the paleoecology of past populations and extinct species of mammals. Most of these studies have focused on terrestrial taxa, including northern hemisphere species such as brown bear (*Ursus arctos*), hyena (*Crocuta crocuta*), musk ox (*Ovibos moschatus*), saiga (*Saiga tatarica*), and woolly rhinoceros (*Coelodonta antiquitatis)* [1–5], and South American species including camelids, Equids, and Gomphotheres (extinct elephant-like proboscideans) [6,7].

To a lesser extent, stable isotope studies have been applied to fossil bone and dentine material of marine mammal species, including pinnipeds and cetaceans [e.g., 8-15]. Stable isotope analysis of osseous materials has also been applied to contemporary marine mammal populations to elucidate foraging ecology and dietary niche of e.g., narwhals (*Monodon monoceros*) and belugas (*Delphinapterus leucas*) [16,17].

As a palaeoecological marker, carbon isotopes provide information on the primary producers at the base of the food web within an ecosystem; carbon reflects their photosynthetic pathways and the environmental parameters affecting them. In terrestrial environments, the primary producers contributing to *δ*^13^C variation are plants [18], while in marine environments, *δ*^13^C primarily reflects phytoplankton and algae composition [12]. For marine mammals and other large predators, gradients in *δ*^13^C also exist such that those with primarily benthic diets tend to have higher tissue *δ*^13^C values than those with primarily pelagic diets [19]. In both terrestrial and marine ecosystems, *δ*^15^N reflects the trophic level of an individual or species [20], however, the *δ*^15^N at the base of the food web also influences the *δ*^15^N of higher trophic level consumers such that a single species, feeding at the same trophic level in two different ocean basins, may have very different tissue *δ*^15^N values [21].

Combined, *δ*^13^C and *δ*^15^N from bone collagen provide information on the average diet of an individual over multiple years [20,22]. In contrast with bone, which is continually remodelled, dentine grows incrementally, and therefore the isotopic composition of dentine corresponds to different years of an individual’s life, depending on the specific tooth/tusk development of the target species [23]. Thus, despite bone and dentine collagen being the same substrate, they incorporate species’ life histories differently. In narwhals, for example, where males have one (and rarely two) long erupted tusks, stable isotope values from the various dentine layers of the tusk represent an individual’s diet in different years [24]. In mammal species, where young nurse on their mother’s milk until they are weaned, collagen extracted from dentine of the first set of teeth in calves/juveniles, or from the roots of the permanent teeth, usually have higher *δ*^15^N values, reflecting the suckling period [2,25,26].

If differences in stable isotope values between bone and dentine are systematic, dentine isotopic values could be translated into the bone-collagen equivalent using a correction factor. This would enable the direct comparison between *δ*^13^C and *δ*^15^N derived from these distinct skeletal elements. The correction factor is an estimate of the average difference between bone and dentine *δ*^13^C and *δ*^15^N from the same individual, and across conspecific samples. This approach has been used to estimate the average difference in *δ*^15^N retrieved from bone and tooth in terrestrial mammals. Two examples of species where correction factors were estimated are wolves and hyenas. For wolves, a correction factor of 1.92 ± 0.19‰ was estimated and, for hyenas, of 1.08 ± 0.55‰ [2]. However, it is important to keep in mind the difference in growth pattern between marine and terrestrial mammals. Marine mammals’ tooth growth pattern is characterized by the layered deposition of dentine, which happens yearly in most marine mammal species [27].

Museum collections housing faunal remains are a crucial resource for studying changes in populations and species across time and space. However, when analysing specimens of the same species from collections or from the field, it is not always possible to access the same skeletal element. Targeting only a specific skeletal element (e.g., cranial bone or tooth) for analysis may drastically reduce the number of available specimens, affecting the robustness and reliability of results. Here, we present a systematic approach to circumvent this issue in two toothed whale species, narwhal and beluga (the only two species within the Monodontidae family).

To calculate a correction factor for *δ*^13^C and *δ*^15^N in narwhal and beluga, we obtained collagen *δ*^13^C and *δ*^15^N from bone and dentine collected from voucher specimens housed at the Natural History Museum of Denmark, University of Copenhagen, and from the Greenland Institute of Natural Resources. The various samples were used to estimate isotopic differences between bone and dentine collagen within individuals, and average differences among individuals within each species. Average differences were used to calculate a correction factor, which can be applied to transform dentine collagen isotopic values into bone-collagen equivalent.

## Materials and Methods

### 1. Samples

The sampled material comprised skulls from 11 narwhals and 26 belugas. We sampled skulls for both bone and dentine at the Natural History Museum of Denmark, University of Copenhagen, and from the Greenland Institute of Natural Resources (see detailed sample information in Supplementary Table 1). Five narwhal skulls were from West Greenland (WG) and six from East Greenland (EG). All belugas were sampled in West Greenland, as the species is not present in East Greenland. The skulls were collected in the field between 1990 and 2018.

Narwhals have a unique dentition; they lack teeth in the lower jaw, and in the upper jaw males have one long erupted tusk (rarely two) and one embedded tooth, while females have two embedded teeth [28]. Hence for narwhals, we determined the isotopic compositions of three skeletal elements: tusk, embedded tooth, and bone. For belugas, which have a row of homodont teeth in both the upper and lower jaw, we sampled tooth and bone.

All individuals sampled for this study were adults, to avoid issues of high *δ*^15^N values in the bone and dentine of nursing calves/juveniles. Sex information for the samples was based on the presence/absence of a tusk in narwhals, and/or the inspection of sexual organs in both narwhals and belugas.

### 2. Stable isotope *δ*^13^C and *δ*^15^N analysis

We drilled 200-300 mg of each skeletal element. For narwhals, we drilled (i) bone, (ii) tusk, and (iii) embedded tooth. For the tusk, we cut a chunk from the base of the tusk. The embedded tooth of narwhal was sampled from well preserved skulls; thus, the embedded teeth were inside the skulls. To sample them, we cut the skulls with a radial drill, creating a hole to access the embedded tooth. Using that hole, we sampled a chunk from the embedded tooth. For one of the narwhal samples (ID: 956), we did not have access to the tusk, and therefore only sampled bone and embedded tooth. For belugas, we drilled tooth dentine from each skull; *δ*^13^C and *δ*^15^N data from the same specimens have been published previously in Skovrind et al. [16] and Louis et al. [17]. In our study, the beluga bone data was used to estimate differences between skeletal elements. See detailed sample information in Supplementary Table 1.

Stable carbon and nitrogen isotope compositions were determined using a continuous flow isotope ratio mass spectrometer at the Water Quality Centre, Trent University, Canada.

Bone and dentine samples were cut into small fragments using an Ultimate XL-D micromotor with a diamond-tipped cutting wheel (NSK-Nakanishi International, Kanuma, Tochigi, Japan) and placed in glass culture tubes. Based on sample availability, some samples were finely powdered while others were left as small chunks as described above. The same pre-treatment protocols described below were applied for both bone and dentine samples.

Each sample was immersed in 2:1 chloroform:methanol solution and sonicated for 1 h to remove lipid contaminants. Following sonication, the sample material was pelleted via centrifugation and the organic solution containing lipids was removed. Another aliquot of the chloroform:methanol solution was then added to each tube for a further hour of sonication. After the second hour of ultrasonication, the solution was removed while leaving the sample pellet intact, and the samples were left to dry for 24 h at room temperature.

Following lipid extraction, the samples were demineralized in strong acid to remove the bioapatite component. To each culture tube, 9 mL of 0.5 M hydrochloric acid (HCl) were added, and samples were placed on an agitating table to increase the surface area of the sample exposed to the acid. The powdered samples had an increased rate of demineralization and were therefore removed from HCl solution after ~6 h. The coarsely-ground samples required longer demineralization periods up to 36 h. Samples were removed from HCl when they appeared translucent or had a malleable texture. Each sample was then rinsed in Type I water (resistivity > 18.2 MΩ·cm) at least 4 times, or until the pH of the solution was 6 as indicated using pH test strips. The samples were then solubilized using 3.5 mL of 0.01 M HCl at 75 °C for 36 h.

Following solubilization, the samples were transferred into glass vials and lyophilized. From a subset of the samples, 0.5 mg of the lyophilized material was weighed into tin capsules for isotopic analysis. The remainder of the samples were then treated using a second lipid extraction protocol to ensure complete removal of lipid contaminants.

The second lipid extraction began with lyophilized material being resuspended in 1.6 mL of Type I water. Next, 6 mL of 2:1 chloroform:methanol was added to the soluble collagen and the samples were then sonicated for 1 h. Following sonication, the samples were centrifuged to generate a three-phase extraction consisting of water-methanol-collagen solution in the top layer of the extraction, with lipid, and chloroform:methanol layers below. The protein-containing top layer of the solution was transferred into a clean tube, and the lipid-chloroform-methanol components of the extraction were left in the tube. An additional aliquot of methanol was then added to the lipid containing portion of the extraction and the same protocol of sonication and centrifugation was followed to try and isolate any protein remaining in the “waste” portion of the extraction. The top layer was again transferred into a clean tube and then left for 24 h at 62 °C to evaporate off the methanol component to isolate the desired water-soluble protein component of the extraction. Finally, an additional 6 mL of 2:1 chloroform:methanol was added to the protein-water portion of the extraction to attempt to remove any remaining lipids. Samples were left to sit for 1 h and then centrifuged. The top layer of the three-phase extraction was removed for the final time and left to evaporate in a new tube for 24 h, then transferred to a glass vial and lyophilized.

Approximately 0.5 mg of lyophilized collagen was weighed into tin capsules and analyzed via a continuous flow isotope ratio mass spectrometer paired with an elemental analyzer. The samples were analyzed using a Euro Vector EA 3000 paired with a Nu Horizon CF-IRMS at the Water Quality Centre, Trent University, Canada. Calibration of sample stable isotope composition was performed relative to the Vienna Pee Dee Belemnite (VPDB) and atmospheric nitrogen (AIR) scales for *δ*^13^C and *δ*^15^N respectively. The calibration was performed using USGS40 and USGS66 or USGS63. In house collagen standards SRM-1 (caribou bone collagen), SRM-2 (walrus bone collagen), and SRM-14 (polar bear bone collagen) were used to monitor accuracy of the stable isotope measurements. Additionally, duplicate samples were interspersed throughout the analyses to measure the homogeneity of the experimental samples relative to known homogenous standards.

Differences in *δ*^13^C in samples collected at different time points may reflect changes in *δ*^13^C of atmospheric CO2, reflecting the industrial revolution and the use of fossil fuels [29]. Therefore, to avoid erroneous interpretations, *δ*^13^C from samples collected across time require correction. We corrected for this change, termed the Suess effect, following the formula from [30].

To contextualise the data generated in our study, we combined our findings with published *δ*^13^C and *δ*^15^N bone records (also from skulls) from 40 WG narwhals, 39 EG narwhals, and one additional beluga [17].

### 3. Skeletal element comparisons

To estimate a correction factor, we calculated pairwise average isotopic differences between skeletal elements for each sampled individual (data for narwhal: Supplementary Table 2; data for beluga: Supplementary Table 3). For narwhal, these estimates were also calculated for each population, West Greenland and East Greenland; results are summarised in Table 1. Correction factors were estimated as the average of the pairwise average differences for each species, and for each population in narwhal.

**Table 1.** Average differences and standard deviations in *δ*^13^C and *δ*^15^N isotopic composition between different skeletal elements in narwhal and beluga. These average differences were used as correction factors to translate dentine values to bone-collagen equivalents.

| Species | Population | Skeletal element | $\Delta\delta^{13}\text{C}$ | $\Delta\delta^{15}\text{N}$ |
| --- | --- | --- | --- | --- |
| | | | Average ± s.d.<br>‰ | Average ± s.d.<br>‰ |
| Narwhal | West Greenland | Tusk/Bone | $0.81 \pm 0.13$ | $0.49 \pm 0.60$ |
| Narwhal | West Greenland | Embedded/Bone | $0.37 \pm 0.45$ | $1.65 \pm 0.69$ |
| Narwhal | East Greenland | Tusk/Bone | $0.31 \pm 0.25$ | $0.48 \pm 0.60$ |
| Narwhal | East Greenland | Embedded/Bone | $0.51 \pm 0.35$ | $1.45 \pm 0.49$ |
| Beluga | West Greenland | Tooth/Bone | $0.37 \pm 0.26$ | $0.59 \pm 0.60$ |

## Results

### 1. Isotopic skeletal differences in narwhals

In narwhals, *δ*^13^C values from bone collagen were lower than in dentine; this pattern was consistent in both WG and EG narwhals. For *δ*^13^C, average values and standard deviations (which will always be reported together with the average values throughout the manuscript) were: bone = −15.43 ± 0.82‰, tusk = −15.08 ± 0.87‰, and embedded tooth= – 14.99 ± 0.75‰. In WG narwhals, differences in *δ*^13^C between tusk and bone (0.81 ± 0.13‰) were larger than differences between embedded tooth and bone (0.37 ± 0.45‰). In EG narwhals, we observed the opposite, and differences between tusk and bone (0.31 ± 0.25‰) were smaller than differences between embedded tooth and bone (0.51 ± 0.35‰) (Table 1, Figure 1).

**Figure 1.**
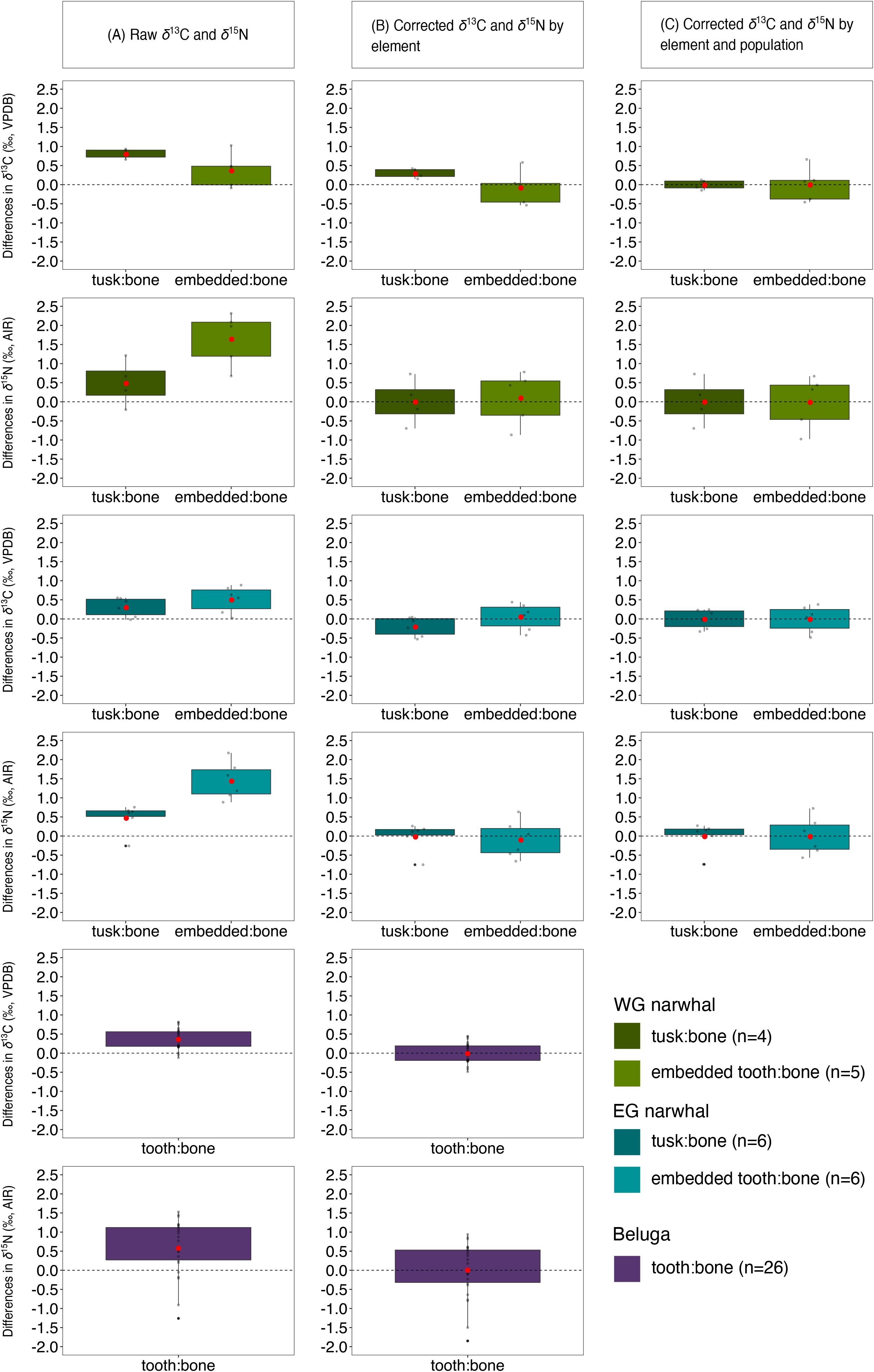
Boxplots showing the average differences in *δ*^13^C and *δ*^15^N composition between various skeletal elements analyzed from narwhal (tusk:bone, embedded tooth:bone) and beluga (tooth:bone). Red dot in each boxplot represents the mean. (A) *δ*^13^C and *δ*^15^N raw composition, with *δ*^13^C corrected for the Suess effect; (B) *δ*^13^C and *δ*^15^N corrected using the correction factor estimated for each species/skeletal element (data for narwhal: Supplementary Table 2; data for beluga: Supplementary Table 3); and (C) *δ*^13^C and *δ*^15^N corrected using the correction factors estimated for each narwhal population, considering known population differences [14]. All data can be found in Supplementary Table 4. Colours represent species and populations. Green palette: West Greenland (WG) narwhal, blue: East Greenland (EG) narwhal, and purple: beluga.

In both narwhal populations, *δ*^15^N values from bone collagen were lower than in dentine. For *δ*^15^N, average values were: bone = 16.14 ± 0.86‰, tusk = 16.54 ± 0.98‰, and embedded tooth= 17.68 ± 1.17‰. Both WG and EG narwhals showed smaller differences between tusk and bone (0.49 ± 0.60‰ and 0.48 ± 0.37‰, respectively), than between embedded tooth and bone (1.65 ± 0.69‰ and 1.45 ± 0.57‰, respectively) (Table 1, Figure 1).

### 2. Isotopic skeletal differences in belugas

In belugas, *δ*^13^C values from bone collagen were lower than in dentine. For *δ*^13^C, average values were: bone = −14.02 ± 0.25‰ and tooth = −13.65 ± 0.23‰. The average *δ*^13^C difference between tooth and bone was 0.37 ± 0.2‰ (Table 1, Figure 1). For *δ*^15^N, average values were: bone = 17.33 ± 0.61‰ and tooth = 17.92 ± 0.63‰. The average *δ*^15^N difference between tooth and bone was 0.59 ± 0.69‰ (Table 1, Figure 1).

### 3. Correction factors in narwhals and belugas

Average differences among skeletal elements for each species and population (for narwhal) were used as a correction factor to translate dentine collagen isotopic values to bone-collagen equivalent (Table 1). Differences in *δ*^13^C and *δ*^15^N before and after correcting the data are visualised in Figure 1. Absolute comparisons can be found in Supplementary Figure 1 and full data can be found in Supplementary Table 4.

We also summarized average *δ*^13^C and *δ*^15^N before and after correcting the data derived from each dentine element (tusk, embedded tooth, tooth); we report these average values for each species, and for the two geographic populations for narwhals in Table 2. Overall, we detected a reduction in the difference between skeletal element average values after application of the correction factor (Table 2, Figure 1). Dentine corrected values fall within the bone average of a combined panel of newly generated and published data for these species (Supplementary Figure 1).

**Table 2.** Average *δ*^13^C and *δ*^15^N values before and after correcting the data for systematic differences in skeletal element composition. WG: West Greenland; EG: East Greenland.

| Species | Population | Skeletal element | Average $\delta^{13}\text{C}$ | | Average $\delta^{15}\text{N}$ | |
| --- | --- | --- | --- | --- | --- | --- |
|  |  |  | Before correcting data (‰) | After correcting data (‰) | Before correcting data (‰) | After correcting data (‰) |
| Narwhal | West Greenland | Tusk | $-14.12 \pm 0.42$ | $-14.93 \pm 0.42$ | $17.44 \pm 0.98$ | $16.95 \pm 0.98$ |
| | | Embedded tooth | $-14.35 \pm 0.33$ | $-14.72 \pm 0.33$ | $18.59 \pm 0.94$ | $16.94 \pm 0.94$ |
| Narwhal | East Greenland | Tusk | $-15.72 \pm 0.18$ | $-16.03 \pm 0.18$ | $15.95 \pm 0.34$ | $15.47 \pm 0.34$ |
| | | Embedded tooth | $-15.52 \pm 0.56$ | $-16.03 \pm 0.56$ | $16.92 \pm 0.69$ | $15.47 \pm 0.69$ |
| Beluga | West Greenland | Tooth | $-13.65 \pm 0.23$ | $-14.02 \pm 0.23$ | $17.92 \pm 0.63$ | $17.33 \pm 0.63$ |

## Discussion and Conclusions

We measured *δ*^13^C and *δ*^15^N from bone and dentine collagen derived from narwhal (n = 11) and beluga (n = 26). The bone and dentine were sampled from whale skulls housed in museum collections. We performed a systematic comparison of isotopic composition of various skeletal elements from the same individuals, and across individuals within each species and/or population (for narwhal). These values were successfully used as a correction factor, allowing the direct comparison of data from different skeletal elements, and from different populations.

For both narwhal and beluga, we detected lower values of *δ*^13^C and *δ*^15^N in bone collagen than in dentine collagen. For narwhal, the highest *δ*^15^N values were found in embedded tooth. Due to the smaller size of the embedded tooth compared to the tusk, its isotopic composition could still present higher *δ*^15^N due to the suckling effect [23,25]. Marine mammal milk is rich in lipids [31] and because lipids have lower *δ*^13^C values relative to proteins [32], we might expect lower *δ*^13^C values in teeth relative to bone collagen [33], but we did not observe this pattern, consistent with the results of several other studies [34–36]. The carbon in collagen is derived primarily from dietary protein [37,38] and the relative importance of lipids in the diet may not be clear when comparing the collagen from bones and teeth. For belugas, Matthews and Ferguson 2015 [36] found consistent declines in the *δ*^15^N values across the first growth layer groups of teeth, consistent with increased *δ*^15^N values during a prolonged period of suckling. They did not find consistent patterns with respect to *δ*^13^C, with about half of the 27 whales they examined showing no variation across the period of nursing inferred from the *δ*^15^N values.

Differences in isotopic composition across tusk layers have been reported [24]; however, our sampling strategy did not target a specific layer, but rather sampled a chunk of tusk comprising several layers, and hence reflected average *δ*^13^C and *δ*^15^N across several years. Because of this sampling strategy, the correction factors generated in this study are applicable to samples collected in a similar manner.

Using *δ*^13^C and *δ*^15^N from bone collagen as dietary indicators, population differences in the foraging ecology of narwhals in West Greenland and in East Greenland have been reported [17]. Our narwhal bone and dentine data also showed systematic differences in *δ*^13^C and *δ*^15^N based on populations. Thus, for narwhal, we estimated a species-wide correction factor, as well as a correction factor specifically for each population, to minimise possible differences in the life history of these two populations that could influence our results (Table 1, Figure 1). When we applied the species-wide correction factor, the average differences between tusk and bone, or embedded tooth and bone, were reduced to values approaching zero (Figure 1B, Figure 1C). Thus, we show that in the absence of *a priori* information on population differences due to life history in isotopic values within a species, it is feasible to apply a species-wide correction factor to combine bone and dentine data. However, if possible, one should consider life history differences among populations of a species when estimating correction factors, as it might bias results.

Our bone/dentine correction factor was estimated based on cranial bone and dentine derived from three dentine elements; tusk and embedded tooth in narwhals, and tooth in belugas. As only one bone element was included (skull), our correction factor may not be fully representative of the intraskeletal bone isotopic variation in either species. *δ*^13^C and *δ*^15^N values in bone tissue are dependent on bone remodeling and biochemical turnover rates [22]. In humans, higher bone turnover rates are significantly and negatively correlated with lower *δ*^15^N [39]. Smith et al. 2020 [40] compiled published and new data on intraskeletal bone variation in *δ*^13^C and *δ*^15^N across marine mammals. The data showed variation in intraskeletal bone *δ*^13^C ranging from 0.1 ‰ in walrus (*Odobenus rosmarus)* (between cranium and mandible [41]) to 1.4 ‰ in dusky dolphin (*Lagenorhynchus obscurus)* (among atlas, humerus and basioccipital [42]). Variation in intraskeletal bone *δ*^15^N ranged from 0.3 ‰ in walrus and dusky dolphin [41,42] to 1.3 ‰ in South American sea lion (*Otaria flavescens*) (among atlas, humerus and basioccipital [42]). However, other studies could not detect intraskeletal variation when performing similar analysis; e.g., [43] did not detect differences in isotopic composition between cortical and trabecular bone in harp seals (*Pagophilus groenlandicus*).

Intraskeletal bone variation in *δ*^13^C and *δ*^15^N has also been investigated in terrestrial mammals (e.g., in humans [39]; in cow (*Bos taurus*), pig (*Sus domesticus*), long-tailed weasel (*Mustela frenata*), and raccoon (*Procyon lotor)* [43]). Using an isotopic distance value estimated from *δ*^13^C and *δ*^15^N bivariate plots, Hyland et al. 2021 [43] detected an average isotopic distance value of bone sampled within individuals of 0.52 ‰. Our results showed intraskeletal variation between bone and various dentine elements. In narwhal, we found a species-wide variation in *δ*^13^C of 0.51 ± 0.33‰ between tusk and bone, and 0.45 ± 0.38 ‰ between embedded tooth and bone. For *δ*^15^N, these values were 0.49 ± 0.44 ‰ and 1.54 ± 0.57 ‰, respectively. In beluga, the *δ*^13^C average difference between tooth and bone was 0.37 ± 0.26 ‰; for *δ*^15^N, it was 0.59 ± 0.69 ‰.

Paleoecological inference from museum specimens is crucial for elucidating changes in populations and species across time and space. However, the same skeletal element is often not available across specimens. Bone and dentine collagen are known to show systematic differences in *δ*^13^C and *δ*^15^N [44]. Thus, analyzing them together without applying a correction factor may influence biological inferences. Here, we estimate and apply a correction factor to our data from narwhal and beluga, and transform dentine values to bone-collagen equivalent. Our approach assumes that one correction factor can be broadly applied to transform dentine collagen to bone-collagen equivalent, and that data from different bone skeletal elements can be analyzed in unison. Application of our correction factors allow the direct comparison of *δ*^13^C and *δ*^15^N retrieved from these different skeletal elements (Figure 1 and Table 2). We acknowledge that our approach may not capture the full variation of skeletal stable isotopes within an individual. However, using the correction factors when analysing stable isotope data retrieved from bone and dentine reduces a large fraction of the intraskeletal variation. This approach has been successfully applied to terrestrial mammals (e.g., [2] reviewed the topic for several species). We suggest applying these correction factors to some kind of bulk dentine sample, rather than to targeted layers of dentine, especially those layers representing the earliest years of life with presumably the highest *δ*^15^N values. In that way, the dentine *δ*^15^N values (as in our case) would be more analogous to bone than those from targeted layers. Here, we show the applicability of skeletal correction factors in two marine mammal species: narwhals and beluga. When dentine values were transformed to bone-collagen equivalents using our approach, they fall within the range of known bone *δ*^13^C and *δ*^15^N values for these species, demonstrating their applicability (Figure 1).

## Acknowledgements

We thank the Institutions that provided us with samples, and the fieldworkers and hunters collecting the samples. We thank Jeppe Møhl and Daniel K. Johansson for assisting with the sampling of the narwhal skulls at the Natural History Museum of Denmark, University of Copenhagen.

## Disclosure statement

The authors report there are no competing interests to declare

## Data Availability Statement

Electronic Supplementary Table 1.

## Author Contributions

Research idea and study design: ARI, PS, EDL; data acquisition: ARI, MK, EG, MPHJ, 294 EDL; data generation: TW, JR, PS; data analysis/interpretation: ARI, TW, JR; manuscript 295 drafting: ARI, EDL with input from all the co-authors.

## Funding

This research was supported by the Villum Fonden Young Investigator Programme under grant no 13151; and the Carlsberg Foundation Distinguished Associate Professor Fellowship under grant no CF16-0202 to EDL.

## Supplementary Figures and Tables

**Supplementary Figure 1.** Boxplots showing *δ*^13^C and *δ*^15^N composition of the various skeletal elements analyzed from narwhal and beluga. Lines in the boxplots represent mean. (A) *δ*^13^C and *δ*^15^N raw composition, with *δ*^13^C corrected for the Suess effect; (B) *δ*^13^C and *δ*^15^N corrected using the correction factor estimated for each species/skeletal element (data for narwhal: Supplementary Table 2; data for beluga: Supplementary Table 3); and (C) *δ*^13^C and *δ*^15^N corrected using the correction factors estimated for each narwhal population, considering known population differences [17]. Colours represent species and /populations. Bone_reference represents the combined newly generated (for this study) and published bone data for beluga and narwhal [16,17]. Green palette: West Greenland (WG) narwhal, blue: East Greenland (EG) narwhal, and purple: beluga.

**Supplementary Table 1.** List of the narwhal and beluga samples used in this study to estimate correction factors.

**Supplementary Table 2.** Correction factor estimates for the narwhal samples.

**Supplementary Table 3.** Correction factor estimates for the beluga samples.

**Supplementary Table 4.** *δ*^13^C and *δ*^15^N data for the narwhal and beluga samples after applying the correction factor estimates. This table also includes the published data (bone_reference) for narwhal and beluga, which was used in Supplementary Figure 1.

## Notes

### Competing Interest Statement

The authors have declared no competing interest.

### Summary of Updates

We have further explained the sampling methods and fully elaborated the stable isotope methods in the main text. We have also highlighted some caveats/recommendations in the discussion

