## Supplementary figures and images for "Combining *δ*^13^C and *δ*^15^N from bone and dentine in marine mammal palaeoecological research: insights from toothed whales"

### Supplementary Figure 1

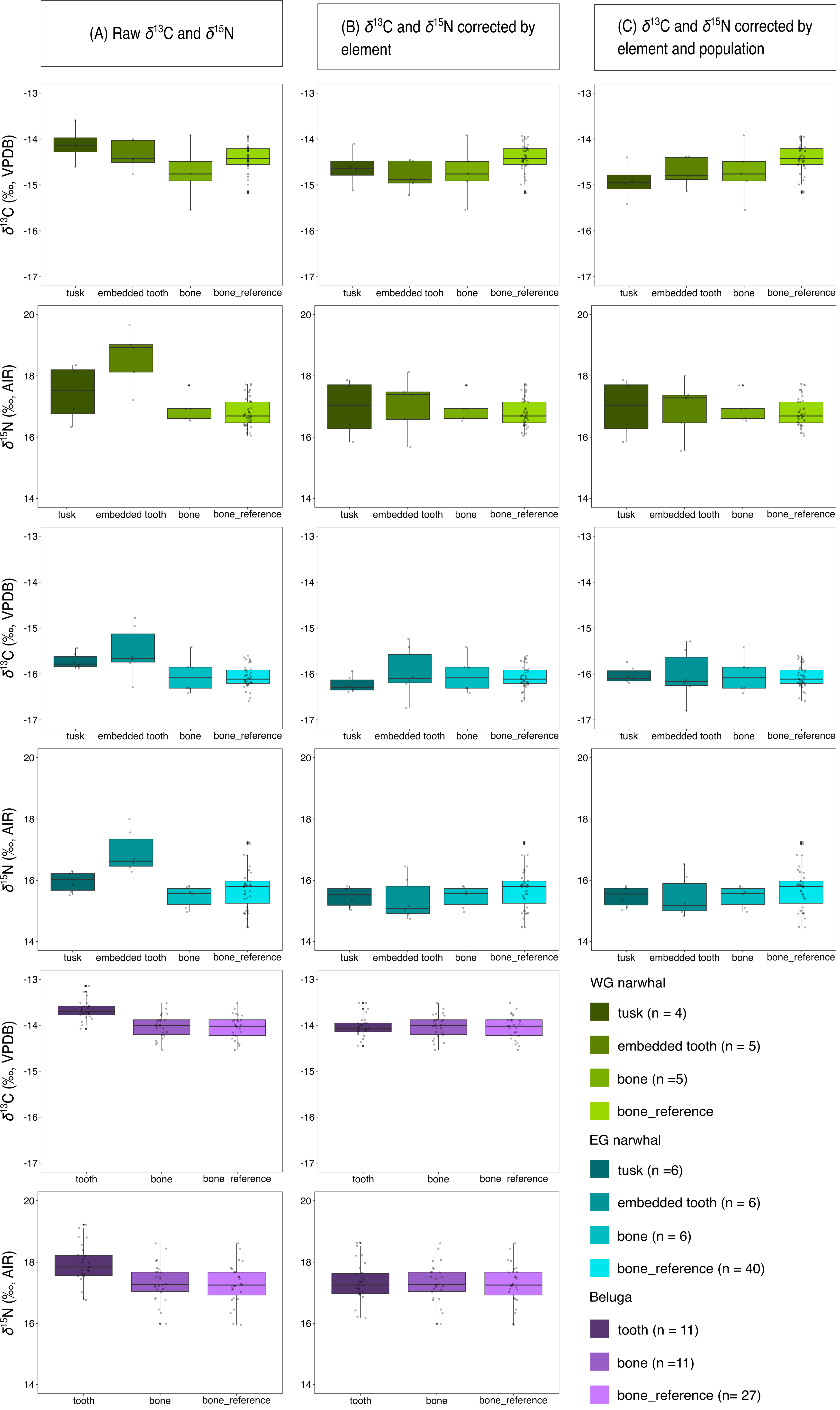
